# The Kinase Chemogenomic Set (KCGS): An open science resource for kinase vulnerability identification

**DOI:** 10.1101/2019.12.22.886523

**Authors:** Carrow I. Wells, Hassan Al-Ali, David M. Andrews, Christopher R. M. Asquith, Alison D. Axtman, Mirra Chung, Ivan Dikic, Daniel Ebner, Jonathan M. Elkins, Peter Ettmayer, Christian Fischer, Mathias Frederiksen, Nathanael S. Gray, Stephanie Hatch, Stefan Knapp, Shudong Lee, Ulrich Lücking, Michael Michaelides, Caitlin E. Mills, Susanne Müller, Dafydd Owen, Alfredo Picado, Kristijan Ramadan, Kumar S. Saikatendu, Martin Schröder, Alexandra Stolz, Mariana Tellechea, Daniel K. Treiber, Brandon J. Turunen, Santiago Vilar, Jinhua Wang, William J. Zuercher, Timothy M. Willson, David H. Drewry

## Abstract

We describe the assembly and annotation of a chemogenomic set of protein kinase inhibitors as an open science resource for studying kinase biology. The set only includes inhibitors that show potent kinase inhibition and a narrow spectrum of activity when screened across a large panel of kinase biochemical assays. Currently, the set contains 187 inhibitors that cover 215 human kinases. The kinase chemogenomic set (KCGS) is the most highly annotated set of selective kinase inhibitors available to researchers for use in cell-based screens.

## Introduction

The protein kinases have emerged as one of the most productive family of drug targets in the 21st century. Over 50 small molecule kinase inhibitors have been approved by the FDA since 2001 for the treatment of cancer, inflammation, and fibrosis [1]. Many of these drugs, specifically those that are used in oncology, owe their efficacy to inhibition of multiple kinases [2]. For some targets second- and third-generation inhibitors have been designed to block the activity of mutant kinases that cause drug resistance after first line therapy. In total these drugs target only a small fraction of the 500+ human kinases. We and others have proposed that the remaining kinases represent an untapped trove of new drug targets [3-5].

Despite the concerted efforts of academic and industrial scientists over the past 25 years, the vast majority of the human kinases remain understudied. Various bibliographic analyses show that, like many other protein families, 90% of the research effort has been expended on < 20% of the kinases [6]. Initiatives such as the NIH-sponsored Illuminating the Druggable Genome (IDG) program have sought to change this dynamic by making available high-quality data sets and research tools for the ‘dark’ kinases [7]. The availability of a set of potent and selective inhibitors of the understudied kinases could greatly aid the study of their biology and uncover new targets for drug development.

An ever-growing number of kinase inhibitors are commercially available. Many of these compounds have advanced to clinical studies and may be useful for investigators seeking to repurpose kinase drugs for a secondary indication but they do little to expand the number of new kinase targets [8]. Notably, the commercially-available kinase inhibitors vary widely in the amount and depth of annotation provided. Their vendors typically list the primary target of each inhibitor and perhaps a handful of off-targets but do not always provide full kinase selectivity profiles. Although some of these data can be found in public databases [9], it is often either unavailable or described in a multitude of assay formats in supporting information accompanying a primary publication. While commercially-available kinase sets contain valuable inhibitors of the well-studied kinases, they do little to provide tools to expand research across the breadth of the kinome.

The public availability of a high-quality chemical probe for every understudied kinase would be an ideal way to embolden researchers to explore the therapeutic potential of each of kinase target [10]. However, the development of potent and selective chemical probes for over 500 kinases would be an insurmountable task using current resources and technologies. A chemogenomic set [11] of kinase inhibitors provides a practical solution to the problem [12]. The vast majority of kinase inhibitors, by virtue of competing with the common cofactor adenosine triphosphate (ATP) in a highly conserved binding site, invariably show some cross-activity on multiple kinases. Landmark studies by Bristol-Myers Squibb and GlaxoSmithKline scientists showed that kinome-wide profiling could identify inhibitors with collateral activity on the understudied kinases [13, 14]. Building on these observations, the kinase inhibitor sets PKIS and PKIS2 were assembled as collections of published kinase inhibitors using the principles that chemical diversity and the inclusion of multiple exemplars of each chemotype would increase the breadth of kinase coverage and aid analysis of phenotypic screening data [15]. Both sets have found widespread use in the research community and demonstrated that repurposed inhibitors from past projects could be used to probe the biology of the kinase they were made to target and their off targets as well. However, the full kinase profile of each inhibitor in PKIS and PKIS2 was not known in advance of their selection, and as a result both sets contained many inhibitors that were either too promiscuous (inhibition of too many kinases) or lacked sufficient target potency to be useful contributors to a chemogenomic set [16]. In spite of this limitation, PKIS and PKIS2 contained many valuable inhibitors for a broad set of understudied kinases. Encouraged by these results we proposed a community experiment to build an optimized kinase chemogenomic set (KCGS) to cover every human kinase [17]. Each inhibitor would have its full kinome profile determined in advance and only those compounds that met a prespecified potency and selectivity would be added to KCGS.

At the outset, we chose to make KCGS an open science experiment. All of the compound structures and associated kinase inhibition and selectivity data would be made publicly available. KCGS would be distributed under a Material Trust Agreement (S1 File) that supported its use as a public resource and prevented the recipients from blocking other researchers from using the set [18]. Eight pharmaceutical companies answered the call to donate kinase inhibitors from their internal compound collections to the effort. Many, but not all of these companies, were current partners in the Structural Genomics Consortium (SGC) [19]. In addition, several leading academic groups contributed compounds to the initiative. To date over 1200 kinase inhibitors have been profiled as candidates for inclusion into KCGS. Here we present the first version of KCGS as well as its initial characterization and examples of its use in cell-based assays. The set will be broadly useful to the scientific community for phenotypic screening to identify the roles of various kinases in biology and disease.

## Results

### Compound selection

Candidate kinase inhibitors were received from GlaxoSmithKline, Pfizer, Takeda, Abbvie, MSD, Bayer, Boehringer Ingelheim, and AstraZeneca. In addition, Vertex gave permission to include their commercially available inhibitors. Academic laboratories that donated inhibitors were Cancer Research UK, Nathanael Gray, and multiple SGC sites. In total 250 new inhibitors were donated to the initiative as candidates to complement the 950 inhibitors of PKIS and PKIS2.

At the outset, we selected the DiscoverX *scan*MAX assay to profile all kinase inhibitors donated to the initiative [20]. The *scan*MAX assay provided kinase binding data on 401 wild type human kinases (Table 1), which was at the time the broadest coverage by any single assay panel [21]. All kinase inhibitors were profiled at a concentration of 1 µM. Using a cut-off of 10% activity remaining (PoC, equivalent to 90% inhibition), an activity profile was determined for each inhibitor and a selectivity index (S_10_) was calculated as the fraction of kinases meeting the cut-off. Compounds with an S_10_ (1 µM) < 0.04 were initially selected for consideration for inclusion in KCGS. For these compounds we performed full-dose response experiments in order to determine K_D_ values for all kinases with PoC < 10% in the *scan*MAX experiment.

**Table 1.** Kinase coverage by KCGS version 1. The human kinases are divided into 10 subfamilies [22]. Kinases: The number of human kinases in each subfamily. Assays: The number of kinases in the DiscoverX *scan*MAX panel. KCGS: The number of kinases covered by an inhibitor. %: The percentage of kinases screened that are covered by an inhibitor.

|  | Kinases | Assays | KCGS | % |
| --- | --- | --- | --- | --- |
| **AGC** | 63 | 46 | 20 | 43 |
| **Atypical** | 34 | 7 | 5 | 71 |
| **CAMK** | 74 | 58 | 28 | 48 |
| **CK1** | 12 | 8 | 3 | 38 |
| **CMGC** | 64 | 60 | 37 | 62 |
| **Lipid** | 20 | 13 | 10 | 77 |
| **Other** | 81 | 51 | 26 | 51 |
| **STE** | 47 | 42 | 13 | 31 |
| **TK** | 90 | 81 | 54 | 67 |
| **TKL** | 43 | 35 | 19 | 54 |
| **Total** | 528 | 401 | 215 | 54 |

For inclusion in KCGS, an inhibitor was selected if the DiscoverX assay panel showed K_D_ < 100 nM on its target kinase and S_10_ (1 µM) < 0.025 in a full panel kinase screen [17]. For inhibitors from PKIS, the assay data from the Nanosyn screening panel of 230 kinases was used to calculate the selectivity index in lieu of submitting the compounds to *scan*MAX. For inhibitors from PKIS2 and the newly donated compounds, the data from the DiscoverX *scan*MAX panel of 401 kinases was used to calculate the selectivity index. Compounds that met the inclusion criteria were manually triaged to maximize the coverage of the human kinome. Our aspirational goal was to include two unique chemotypes for each kinase and care was taken not to over represent kinases that had been more heavily studied (such as EGFR, MAPK14, GSK3, and ERBB2). For the poorly studied dark kinases, there was often only one or two compounds to select that met the inclusion criteria. Finally, in those cases where two compounds had equivalent kinase profiles, preference was given to inclusion of the chemotype with fewer exemplars in the set. Using these guidelines, version 1.0 of KCGS was assembled with a total of 187 kinase inhibitors. Summary information for each inhibitor is contained in S1 Table and the full kinase profiles can be accessed at www.randomactsofkinase.org.

### Kinase coverage

The set covered a total of 215 human kinases, which was more than 50% of the full *scan*MAX assay panel (Table 1 and S2 Table). Across the branches of the kinome, the broad coverage was found in the TK family (67%) and CMGC family (62%). While KCGS appears to cover 71% of the atypical kinases, only small fraction of these kinases has assays in the *scan*MAX panel. Lower coverage was seen for the CK1 (38%) and STE (31%) families. 114 kinases were covered by two or more inhibitors, while the remaining 98 kinases have only one useful inhibitor in the current set (S1 Table). Ideally every kinase would be covered by inhibitors from multiple chemotypes to aid analysis of phenotypic screening data and this remains a goal for future expansion of the set.

Despite the tractability of kinases as drug targets, the majority of the kinome is poorly annotated and remains dark with respect to its role in human biology, in part due to a paucity of reagents. The NIH IDG initiative has nominated 162 dark kinases (Fig 1 and S2 Table) for development of chemical and biological tools in an effort to seed research on these understudied proteins [7]. KCGS contain inhibitors of 37 of the IDG dark kinases (S2 Table), which may be useful as initial chemical tools to study these kinases. These KCGS compounds can also be used as starting points for development of high-quality chemical probes as the biological function of their kinase targets becomes better understood and the investment in additional optimization is warranted. We utilized a recent data set [23, 24] annotated to human protein-coding genes and their genetic relevance to look at the frequency of PubMed publications on individual kinases. Fig 1 depicts the publication counts for kinases covered by KCGS. The set contains inhibitors of most of the more highly studied (top 25%) kinases, but importantly it also contains inhibitors that cover some of the darkest kinases. Future expansion of KCGS will focus on filling the gaps in coverage of the dark kinases.

**Fig 1.**
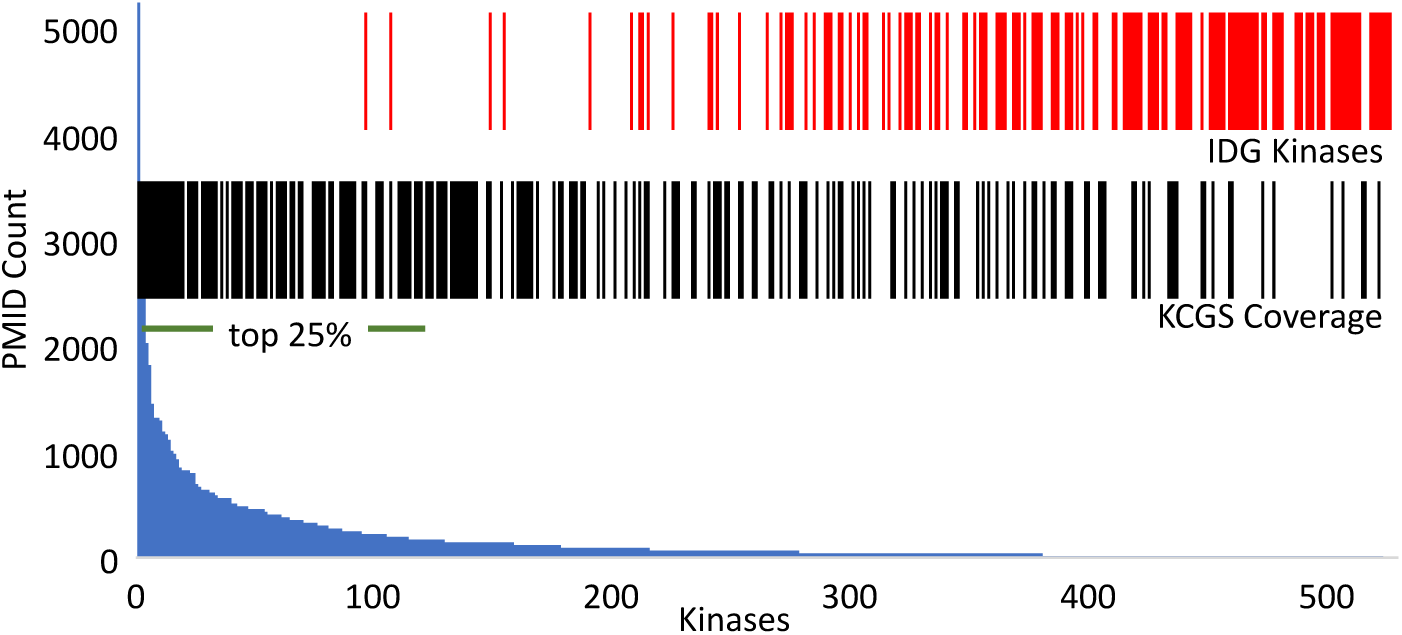
Analysis of kinase coverage in KCGS by publication frequency. The blue histogram indicates the number of publications for each kinase in PubMed (Table S2) [23, 24] ordered by frequency with the top 25% noted by the horizontal bar. The red vertical bars indicate the 162 poorly studied dark kinases nominated by the NIH IDG initiative [7]. The black vertical bars indicate the 215 kinases covered by an inhibitor in KCGS version 1.

### Chemotype analysis

To aid analysis of screening data and to support future expansion of KCGS, a method was developed to assign each inhibitor to a specific chemotype based on the chemical structure of its hinge-binding moiety. To accomplish this, 119 known kinase hinge-binder substructures were defined manually and codified using SMILES arbitrary target specification (SMARTS) [25]. To resolve issues where an inhibitor could be assigned to multiple bins, SMARTS were given a priority order. Kinase inhibitors that lacked an obvious hinge binding group were grouped separately into an additional SMARTS bin. Applying this analysis to the 187 KCGS inhibitors, the compounds were found to occupy 67 of the 120 SMARTS bins (Fig 2A and S3 Table). Nine of the bins contained six or more inhibitors, 27 bins had two to five members, and 31 bins contained only one exemplar. The nine most highly populated SMARTS bins contain well known kinase inhibitor scaffolds, such as indazoles, oxindoles, quinazolines, quinolines, and pyrimidines. Six KCGS compounds that lack an obvious hinge-binder group were placed in the “other” bin. They include an allosteric PAK inhibitor and two allosteric MEK inhibitors. For bins containing multiple exemplars, the individual inhibitors often showed activity on kinases located in several different branches of the kinome. For example, the 13 oxindoles in KCGS showed a cluster of activity on CMGC kinases, but they also inhibited TK, TKL, and STE kinase (Fig 2B). While the oxindole chemotype has been found in many highly promiscuous kinase inhibitors, the inclusion of several oxindoles in KCGS demonstrated that this chemotype can also produce highly selective kinase inhibitors by judicious optimization of the molecules. KCGS contained 9 exemplars in the 4-anilino-quinazolines bin (Fig 2C). Six FDA-approved kinase inhibitors that target EGFR and ERBB2 also fall into this bin. However, the SMARTS analysis highlights that modification of the 4-anilino-quinazoline chemotype can also generate inhibitors with activity on several adjacent kinase subfamilies.

**Fig 2.**
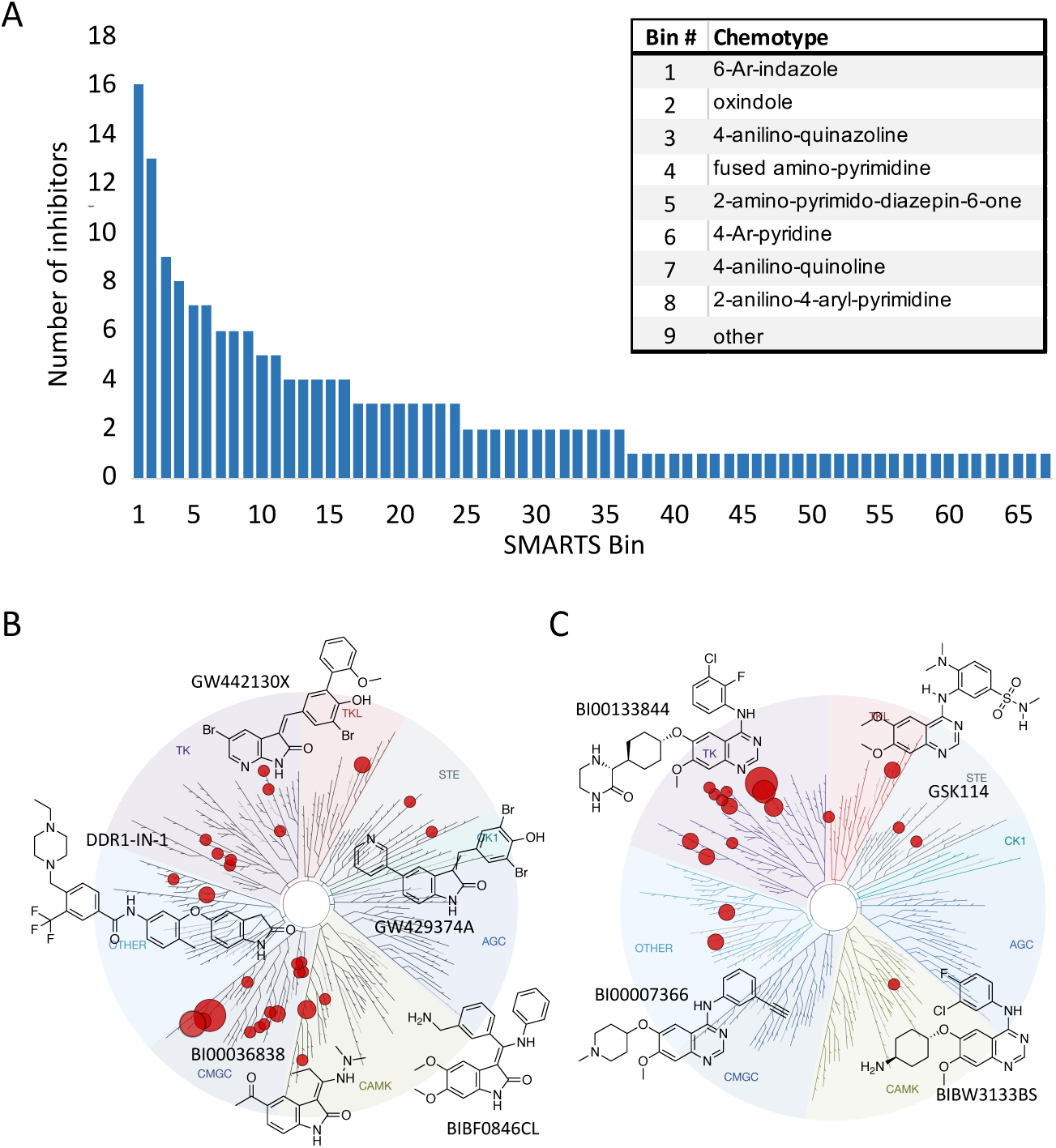
KCGS contains inhibitors from 67 distinct chemotypes. (**A**) The dendrogram displays the number of inhibitors in each SMARTS bin. The inset names the 9 most highly populated bins. (**B**) Tree plot of the human kinases with each subfamily uniquely shaded. Kinases covered by a member of the oxindole SMARTS bin are displayed as red dots, scaled by the number of compounds inhibiting a specific kinase. Representative chemical structures form the oxidole SMARTS bin are shown. (**C**) Kinases covered by the 4-anilino-quinazoline SMARTS bin with representative chemical structures.

### Calculated properties

All of the inhibitors in KCGS were originally the product of medicinal chemistry projects to target specific kinases. As such many of them had been optimized with an eye on physicochemical properties and cellular activity. To evaluate the overall quality of the set, the calculated properties of each inhibitor from KCGS were determined in SwissADME [26] and compared to a set of 52 FDA-approved kinase inhibitors [27]. SMILES strings representing each inhibitor were input to SwissADME to generate the predicted solubility and calculated lipophilicity (S1 Table). For predicted solubility the inhibitors were binned into four categories ranging from poorly to very soluble. The solubility profile of both inhibitor sets was similar (Fig 3A). The majority of compounds were predicted to be moderately soluble or better for both KCGS (88%) and the FDA-approved inhibitors (78%). LogP is a common measure of lipophilicity and is considered a critical factor in assessing the drug-like properties of small molecules [28]. The SwissADME consensus logP (cLogP), which is the arithmetic mean of five calculated values (XLOGP3, WLOGP, MLOGP, SILICOS-IT, and iLOGP), was used to compare KCGS to the FDA-approved kinase inhibitors. The results showed that KCGS clogP values trended towards lower lipophilicity than the FDA-approved drugs, with 65% of the KCGS inhibitors falling between cLogP 2–4 and proportionally fewer inhibitors with cLogP > 4 (Fig 3B). Overall these calculations support the premise that the inhibitors in KCGS have physical properties and are well suited for use in cell-based assays.

**Fig 3.**
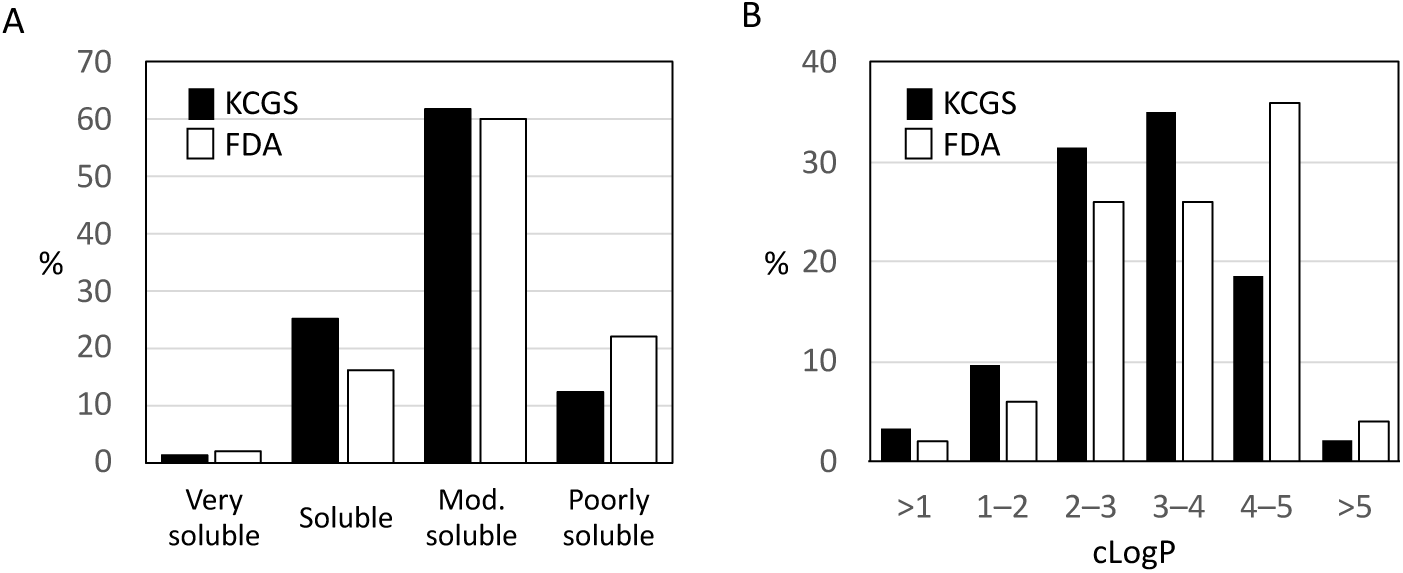
Calculated properties. **(A)** Predicted solubility of the KCGS compounds (black) compared to 52 FDA-approved kinase inhibitors (white) split into four categories and shown as percentage of the set. (**B**) Hydrophobicity analysis of the KCGS compounds (black) using SwissADME compared to the FDA-approved kinase inhibitors (white) split into six ranges of cLogP and shown as percentage of the set.

### Chemogenomic screening

To format KCGS for distribution to a large number of researchers, 10 mM DMSO stock solutions of the 187 inhibitors were aliquoted into 384-well format plates. 1 µL of each inhibitor was dispensed to each well of the plate using an Echo 550 acoustic dispenser for accurate delivery. This 1 µL/10 mM volume provides sufficient compound to run 100 assays at a 1 µM inhibitor concentration in 96-well format and 200 assays in a 384-well format (assuming 100 µL and 50 µL working volume respectively, see S2 File). KCGS is delivered with a plate map that delineates compound identification numbers as well as the kinase profile for each compound (S1 Table). Based on the kinase selectivity profile of the inhibitors, 1 µM is the recommended screening concentration for chemogenomic experiments to support hit identification and target deconvolution. Screening at higher concentrations is likely to complicate data interpretation due to additional undocumented off-target activity of the inhibitors. To aid with hit follow-up additional quantities of each inhibitor are available from the SGC-UNC for full-dose response and secondary assays.

### Cell toxicity

To facilitate the use of KCGS in cell-based assays, we determined the acute toxicity of the individual inhibitors at a high dose (10 µM) in HeLa cells. After 24 h treatment, high content imaging [29] was used to measure healthy cell count as well as the percent necrotic and apoptotic cells, which identified those inhibitors that exhibited varying degrees of toxicity (Fig 4 and S4 Table). 134 of the kinase inhibitors had little or no effect on total cell count. 43 of the inhibitors reduced cell count by 20% or more. The most toxic compounds that decreased cell count by >67% are highlighted in Figure 4A. The cell toxicity displayed by these compounds may be due to their inhibition of kinases that affect cell division or cell viability, either as a primary or secondary target. Among the compounds with the largest effect on cell count were inhibitors of kinases involved in cell cycle progression, checkpoint regulation, and cell division including the CDKs (GW416981X, THZ531, BI00036838), AURKC (GW814408X), ATR (VE-822), and CHK1/2 (CCT244747, CCT241533). Others compounds with significant toxicity in HeLa cells at the 10 µM dose were GW683134A, a type II inhibitor of KDR, KIT and TEK, and PFE-PKIS 29, a very potent (< 10 nM) inhibitor of mTOR and several lipid kinases including all isoforms of PI3K.

**Fig 4.**
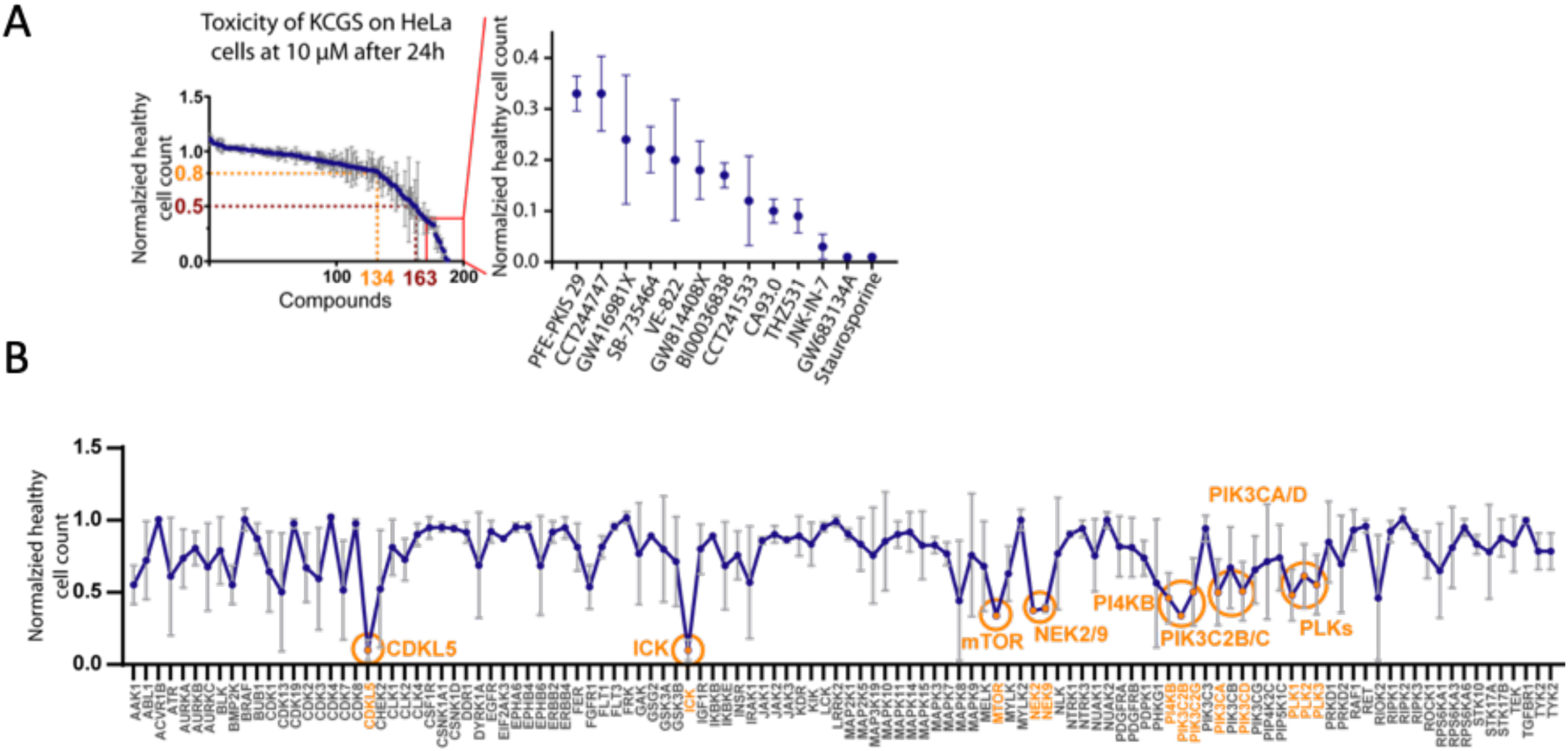
Cell toxicity assessment. (**A**) Effects of KCGS compounds at 10 µM on HeLa cells after 24 h. Measurements were made in triplicate and with standard errors shown. Shown is the normalized healthy cell count with highlighted thresholds of 0.8 (80% healthy cells) and 0.5 (50% healthy cells). The panel on the right side displays the compounds with the greatest effect on healthy cell count in HeLa cells. (**B**) Averaged toxicity measured by normalized healthy cell count for every target covered by two or more chemotypes. Highlighted are target kinases that show a significantly lower healthy cell count than the DMSO control.

To gain further insight into kinases associated with cell toxicity, we performed an analysis of every kinase that is covered in KCGS by two or more distinct chemotypes (Fig 4B). Several kinases were identified whose inhibition resulted in significantly lower healthy cell count. Several inhibitors of the polo-like kinases (PLK) showed toxicity, as did inhibitors of the PIK3C and PI4KB lipid kinases. The apparent toxicity of NEK2/NEK9 inhibition by GSK579289A, GSK461364, GSK579289, and GSK237701A may also be attributed to the collateral PLK inhibition of these compounds at the high concentration that the assay was performed. The apparent toxicity of inhibition of dark kinases CDKL5 and ICK by JNK-IN-7 and BI00036838 may also be due to the inhibition of the other kinase targets of these two inhibitors (S1 Table).

### Cell growth

To further document the effect of KGCS on cell viability we performed assays for cell growth [30] in 16 immortalized cell lines that were selected to cover breast, ovarian, prostate, colorectal, lung, skin, brain, and pancreatic cancer (S5 Table). Non-malignant breast and lung cell lines were included for comparison. Using a 1 µL aliquot of KCGS (10 mM in DMSO), the set was screened in duplicate across the 18 cell lines at a compound concentration of 1 µM. The effects on cell growth and viability were determined after 72 h treatment using high-throughput microscopy [31]. Growth rate inhibition metrics (GR), employed to account for variable division times, were computed [32]. One cell line (SW1783) did not grow under the assay conditions and was excluded from analyses (S6 Table). The effect of KCGS across the remaining 17 cell lines is depicted in Fig 5.

**Fig 5.**
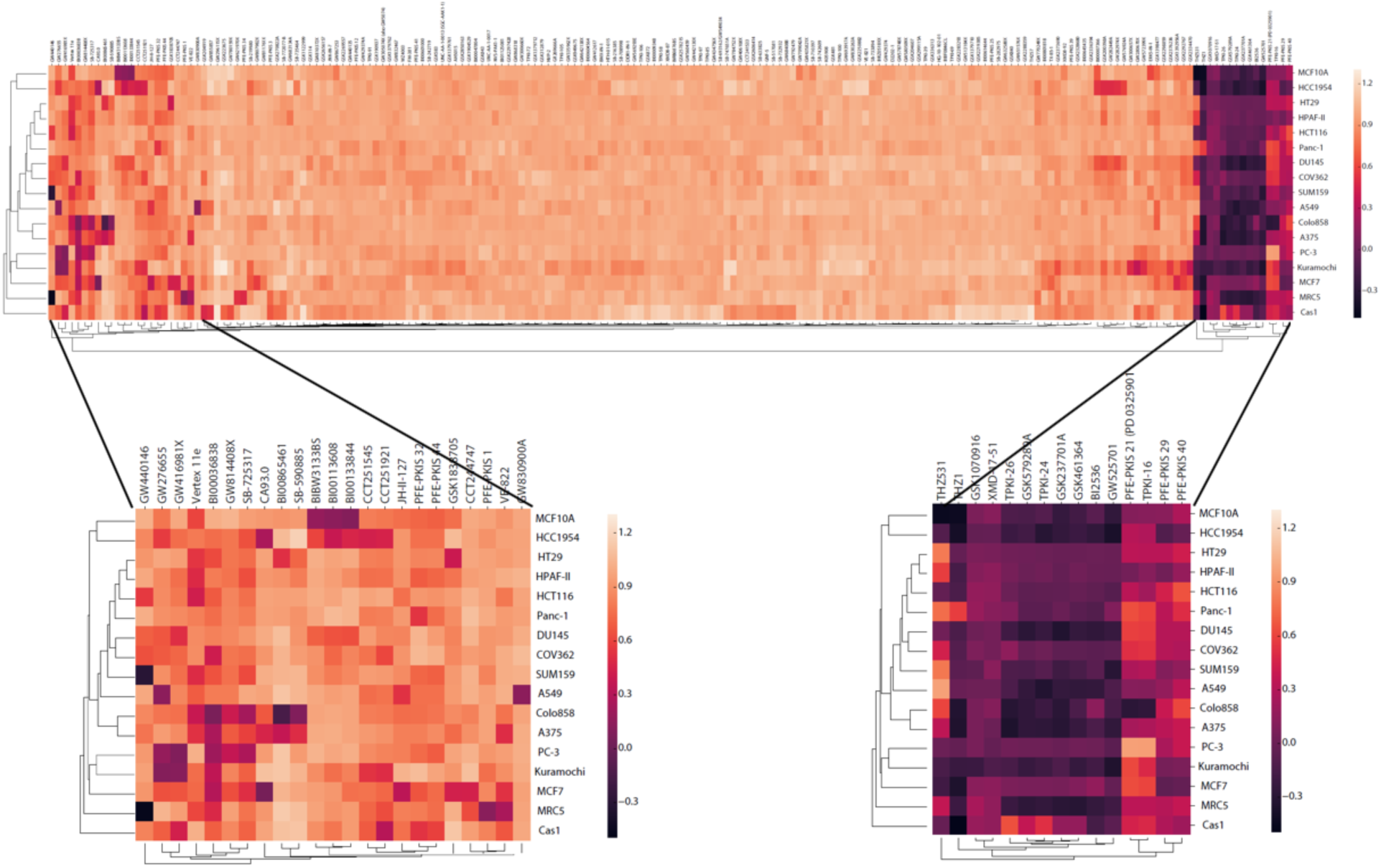
The kinase inhibitors in KCGS have different effects on cell growth across a wide range of lines. Growth rate (GR) was determined at 72 h post compound treatment. Each row displays data from a single cell line. Each column displays data from a single compound. GR is colored from −0.5 in black to 1.3 in yellow. The expanded views display data for compounds that showed cell-line selective decrease in GR and relatively cell-line independent effects, respectively.

Analysis of GR across the 17 cell lines identified three categories of kinase inhibitors. The first category included the majority of inhibitors in KCGS that showed no discernable effect on cell growth, with GR values within 10% of the DMSO control. The second category containing 15 inhibitors showed > 30% decrease in GR across most of the cell lines. Six of these compounds (TPKI-24, TPKI-26, GSK461364, GSK579289A, GSK237701A, and BI2536) have PLK inhibitory activity, two are allosteric MEK inhibitors (PFE-PKIS 21 and TPKI-16), and two are inhibitors of aurora kinases (XMD-17-51 and GSK1070916). Notably, only two of these inhibitors (THZ531 and PFE-PKIS 29) were shown to be cytotoxic at a higher concentration in the HeLa cell experiment. The third category contained 23 inhibitors that showed cell line-dependent effects on GR. Six of these compounds (GW416981, BI00036838, GW814408X, SGK-GAK-1 (CA93.0), CCT244747, and VE-882) had been identified as toxic to HeLa cells, but when tested at 1 µM across a wider range of lines their effects were now shown to be dependent on other cellular factors and not intrinsic to the compounds alone. Based on the annotation of these compounds, inhibition of several kinases was highlighted as being responsible for the cell line-dependent effects. These kinases include multiple CDK isozymes, GAK, BRAF, and BLK. Determination of whether selective inhibition of these kinases would have potential therapeutic utility in specific cancers will require confirmatory follow-up studies such as CRISPR dropout screens and screening of alternate inhibitors of the same targets. However, these data highlight the power of screening a chemically and biologically diverse chemogenomic set of kinase inhibitors to determine how they perturb a simple cell phenotype.

### Kinases linked to autophagy

Autophagy is a central mechanism that helps maintain cellular homeostasis. Autophagy is activated in response to different stress conditions such as starvation, protein aggregation, oxidative stress, bacterial infection, inhibition of the TOR1 pathway, and others [33, 34]. To determine the effect of kinases on autophagic flux, the KCGS library was screened at 1 µM concentration in RPE1 cells stably expressing the autophagic flux reporter construct GFP-LC3B-RFP-LC3BΔC [35] (Fig 6). The cells were monitored in a time-dependent manner so that the ratio of GFP/RFP intensity ratio represented the level of autophagic flux. The averaged GFP/RFP ratio was subsequently normalized to time point 0 h in order to facilitate easy visualization of differences to the autophagy control compounds Torin1 (inducer) and Torin1+ Bafilomycin A (inhibitor and deacidifier of lysosomes) compared to the DMSO vehicle control (S7 Table).

**Fig 6.**
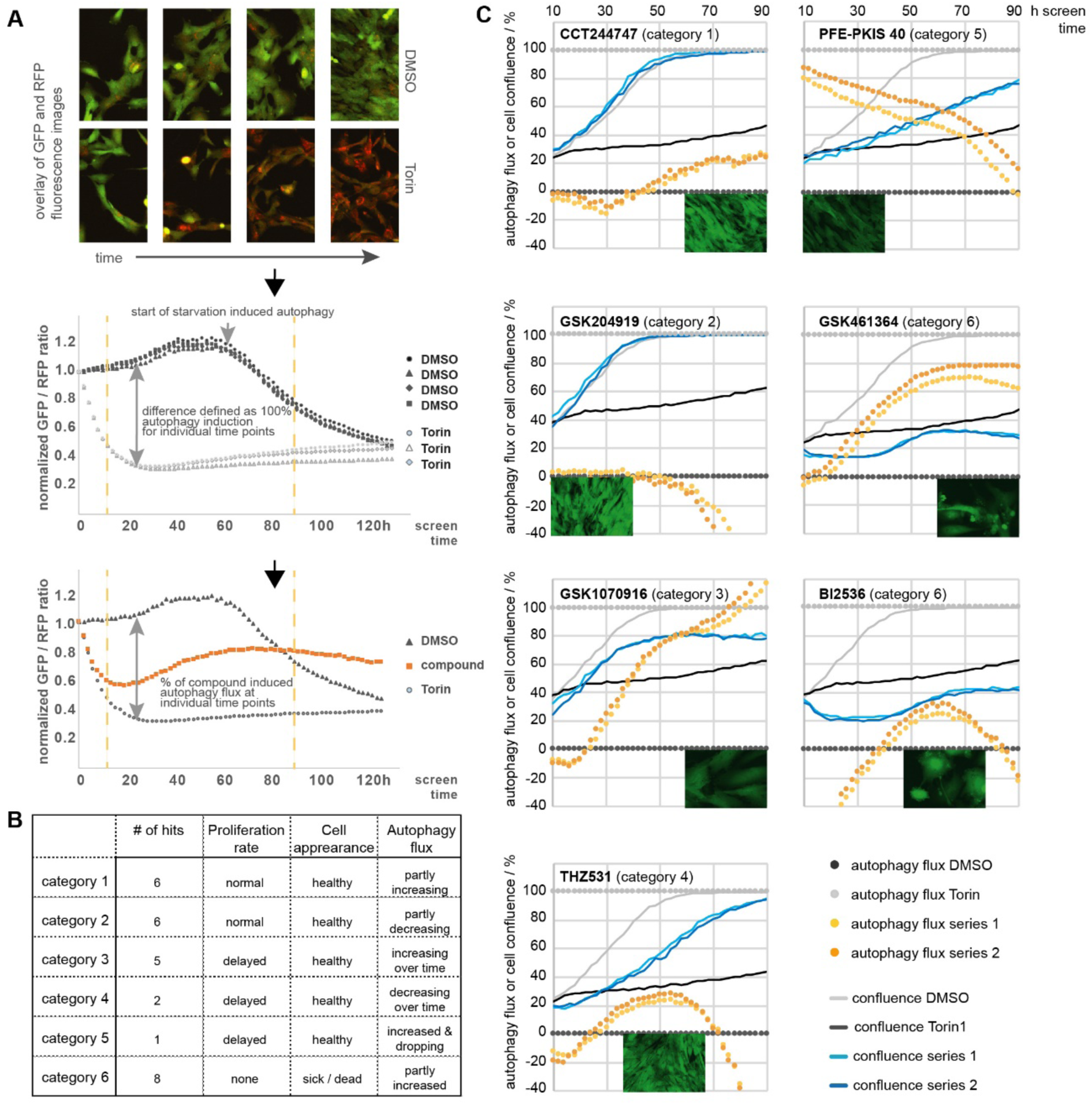
Autophagic flux assay. Compounds (1 µM) were analyzed for their effect on autophagy flux in RPE1 cells stably expressing the general autophagy flux reporter GFP-LC3B-RFP-LC3ΔC. Phase contrast as well as fluorescent images (GFP, RFP) were taken at indicated time points. (**A**) Ratio of the GFP/RFP signal correlating with high (Torin1; defined as 100%) and low (DMSO; defined as 0%) autophagy flux. Compound-induced changes in autophagic flux are represented as percentage of this difference. (**B**) Hits are defined as compounds showing > 20% aberration of the GFP/RFP ratio compared in five or more consecutive time points and categorized according to their cell proliferation rate as well as visual appearance. (**C**) Examples of compounds with an effect on autophagic flux. Cell confluence is presented as percentage of covered growth area. Visual appearance of cells and cell confluence were ascertained by examining fluorescent images of the GFP channel at 96 h screen time. Graphs contain the individual average of experiments run in triplicate at two different times.

As expected, DMSO-treated cells had a stable level of autophagic flux until nutrients in the media were depleted and they entered a starvation induced phase of autophagy induction. In contrast, the GFP/RFP ratio of Torin1-treated cells rapidly decreased within the first screening hours and stayed low over the complete time period of the five-day screen. Of note, treatment with Torin I caused an arrest of cell proliferation but no cell death. Confluence analysis was based on phase contrast images corresponding to the cell-covered area of each well. In this assay, cell health can only be assessed based on visual appearance of the cells and cell proliferation.

Hits were defined as compounds that showed > 20% aberration of the GFP/RFP ratio in at least 5 or more consecutive time points, equivalent to a 10 h assay window. Hits were grouped in 6 categories based on the increase or decrease in proliferation rate, cell appearance and autophagy flux, respectively (detailed in Fig 6). Category 1 hits included GW416981X, a potent CDK1–3 inhibitor. The CDK inhibitors roscovitine and purvalanol have previously been shown to induce autophagy [36]. Likewise, CHK1 inhibition, represented by the category 1 hit CCT244747, has been previously linked to autophagy induction [37]. However, we also identified several new potential kinase targets for autophagy. For example, GSK204919, a potent dual PRKD1/2 inhibitor, caused a reduction in autophagic flux. The role of PRKD in autophagy has not been well-documented. Additional studies are required to link the observed autophagy reduction to PRKD inhibition rather than another kinase target of GSK204919 (e.g. JAK). Categories 3 and 5 compounds contained inhibitors of kinases known to induce autophagy, such as GSK1070916 (an aurora kinase [38] inhibitor) and PFE-PKIS 40 (a PI3K and mTOR [39] inhibitor. Notably, the RPE-1 cells used in the screen did not show any reduction of cell proliferation at 1 µM PFE-PKIS 40, despite its toxicity at 10 µM in HeLa cells (see above). The behavior of the category 4 compounds THZ531 (CDK inhibitor) and PFE-PKIS 29 (mTOR inhibitor) likely resulted from overlap of autophagy induction with cell toxicity as already identified in the cell health and cell growth assays. Most of the compounds in category 6 have also been identified in the cell growth assays, including several inhibitors with activity on PLK (TPKI-24, TPKI-26, GSK461364, GSK579289A, GSK237701A, and BI2536), a kinase known to regulate both autophagy as well as mitosis [40].

## Discussion

KCGS version 1.0 is currently the best publicly available set of well-annotated potent and selective kinase inhibitors. All of the inhibitors have narrow selectivity profiles as ascertained from screening across an assay panel covering the majority of the human protein kinases. The set can be obtained by any investigator who agrees to the open science principles of not restricting its use by others and also promises to publish the results of their screen (S1 File). This manuscript describes the chemical structure and kinase annotation of all of the inhibitors in the current set. We recognize that there is additional room for improvement in the breadth (more kinases) and depth (more chemotypes per kinase) of kinase coverage and in the biological annotation of the set. However, initial characterization of KCGS in phenotypic screens confirmed the utility of the set for chemogenomic exploration of kinase signaling. Screening across 18 cell lines identified a subset of compounds that selectively inhibit their growth. Some of these compounds point to dark kinases that have received little attention as potential drug targets. A screen for autophagy uncovered additional kinase pathways that warrant for further exploration. The narrow spectrum kinase activity of the individual inhibitors and the accompanying annotation supports rapid identification of target kinases for additional study. While, the compounds are generally non-toxic, we recommend that KGCS is screened at a maximum concentration of 1 µM in cells to minimize the potential for inhibition of additional kinases or off-target toxicity.

Several ongoing activities will support KCGS remaining the best publicly available set of kinase inhibitors. Currently, 51% of the screenable kinome, as defined by the DiscoverX *scan*MAX, is covered by KCGS version 1.0 for a total of 215 human kinases. Over 100 of these kinases were selectively inhibited by two or more chemotypes in the set. Our originally stated goal was to cover all human kinases with two or more chemotypes, so additional inhibitors are still required for those kinases that are covered by only single chemotype. There are an additional 250 ‘gap kinases’ where we are still seeking an inhibitor that meets our minimal potency and selectivity criteria for inclusion in the set. For many of the gap kinases that are routinely screened in the DiscoverX *scan*MAX, identification of a non-promiscuous inhibitor is the primary challenge but may be achievable through iterative medicinal chemistry to improve selectivity. Additionally, there are over 50 human kinases for which robust biochemical screening assays are not readily accessible in any format. For these kinases, it is not yet known if useful inhibitors already exist in the current set or among molecules that are in the public domain. Many of these dark kinases are difficult to express and purify or represent pseudokinases with little or no catalytic activity. Development of new screening formats or assay methodologies will be required to identify a complete set of inhibitors for the whole kinome.

One limitation to the design and selection of the inhibitors in KCGS was the use of potency and selectivity data from cell-free biochemical assays. The activity of kinase inhibitors in cells can be affect the binding of small molecules to the kinase. In addition, some inhibitors may be less potent in cells if they are not efficient at crossing the cell membrane. However, the provenance of compounds included in KCGS, either from lead optimization programs in the pharmaceutical industry or the product of academic chemical probe development projects, suggests that most of them are likely to be cell active. In fact, the profile of physical properties across KCGS is as good if not better than a set of 52 FDA-approved kinase inhibitor drugs. Regardless, it is not uncommon for kinase inhibitors to demonstrate lower potency in cells than in cell-free assays. A recent advance in application of NanoBRET technology to measure the target engagement of kinase inhibitors in living cells has provided a method to study this issue [41]. NanoBRET assays have now been developed for 133 of the human kinases that are inhibited by the molecules in KCGS version 1.0. Using these NanoBRET assays we have begun the process of annotating each of the inhibitors in KCGS for its activity against its corresponding target kinase in live cells. These data will aid the deconvolution of phenotypic screening data of KCGS and identify kinases for which inhibitors with improved cell activity will be required in future releases.

The inhibitors in KCGS were sourced from multiple industrial and academic laboratories in a conscientious effort to maximize both the number of chemotypes (chemical diversity) and the breadth of kinase coverage (biological diversity). While the core of the set is still composed of molecules that were published by GlaxoSmithKline chemists, KCGS version 1.0 contains inhibitors that originated from the laboratories of four pharmaceutical companies and three academic laboratories. We continue to seek new inhibitors to add to KCGS that represent either a new chemotype or an inhibitor of a gap kinase. To this end, we have completed profiling of molecules that have been donated by three additional companies as well as molecules synthesized in our laboratories and by academic collaborators. Inhibitors representing new chemotypes that increase the depth of coverage on many kinases will be made available as a supplemental set (KCGS version 1.1). Identification of potent and selective inhibitors of the gap kinases represents a more formidable yet surmountable challenge. We continue to welcome donations of candidate inhibitors of these kinases from industrial and academic laboratories to support expansion of the KCGS. All donor laboratories receive in return copies of the full KCGS set and the satisfaction of contributing to the goal of maintaining the best publicly available set of a high-quality kinase inhibitors.

## Conclusion

KCGS is the largest fully annotated set of selective kinase inhibitors that is accessible to the scientific community. It is available in a format to support phenotypic screening in cells. Importantly, the set is a key resource to support expansion of the druggable genome. Biological annotation of a common set of diverse kinase inhibitors will deepen our understanding of the role of kinases in cell signaling and may uncover new targets for drug discovery programs and precursors to new medicines.

## Materials and Methods

### KCGS

The current version of KCGS is available in 1 µL aliquots of a 10 mM DMSO solution at www.sgc-unc.org/request-kcgs/

### Kinase assays

Compounds were screened at 1 µM using the KINOMEscan technology in the *scan*MAX assay panel of 401 wild-type human kinase and S_10_ was calculated as previously described [17]. Compounds with S_10_ < 0.04 were submitted for K_D_ measurement on kinases with POC < 20%. Compounds with K_D_ < 100 nM and S_10_ (1 µM) < 0.025 were selected as candidates for inclusion in KCGS.

### Chemotype binning

Each molecular substructure or bin representing the desired hinge binder was manually codified in SMARTS language [25] and were given a priority order. Each molecule in KCGS was represented as a SMILES code. The SMARTS search was performed using Open Babel [42] to generate a .smi file to associate the SMILES code of each molecule with a specific SMARTS bin. All .smi files were processed in MATLAB [43] to create a compound-SMART matrix. Compounds with multiple matches were assigned to bin that corresponded to the highest priority SMARTS.

### Cytotoxicity

A triple staining high content screen was performed as previously described [29]. HeLa cells exposed to KCGS compounds at 10 µM for 24 h were stained with the three dyes: Hoechst 33342 (1 µM), Yo-Pro 3 (1 µM) and Annexin V (0.3 µL per well) for 1 h. Cellular fluorescence was measured using the JQ1 high content imaging system (Yokogawa) with the following setup parameters: Brightfield transmitted light at 70% for 50 ms; Hoechst 33342 was excited by 100 ms exposure; Ex 405 nm/Em 447/60 nm, Yo-Pro 3 by 100 ms exposure; Ex 561 nm/Em 617/73 nm and Annexin V (Alexa 488) by 50 ms exposure; Ex 488 nm/Em 525/50 nm. All data were analyzed by the Pathfinder software and four categories for cells were designated: healthy cells, early apoptosis, late apoptosis, and necrosis. Each category was calculated as percentage for every inhibitor. Cell nuclei were classified as either healthy, pyknosed, or fragmented.

### Cell Growth

The KCGS library compounds were arrayed in a 384-well plate at a concentration of 1 mM. 4 breast cell lines, (SUM159, MCF7, MCF10A, and HCC1954), 2 ovarian cell lines (COV362 and KURAMOCHI), 2 prostate cell line (PC-3 and DU145), 2 colorectal cell lines (HT29 and HCT116), 2 lung cell lines (A549 and MRC-5), 2 melanoma cell lines (COLO858 and A375), 2 glioblastoma cell lines (Cas1 and SW1783), and 2 pancreatic cell lines (Panc-1 and HPAF-II) were maintained in their recommended growth media at 37 °C in 5% CO_2_, were seeded in 384-well CellCarrier plates (Perkin Elmer) at the densities listed in S5 Table. Cells were allowed to adhere for 24 hours and treated in duplicate with the KCGS library by pin transfer for a final concentration of 1 µM. Cells were stained and fixed at the time of pin transfer and following 72 hours of treatment. Cells were pulsed for one hour with EdU (Lumiprobe, Hunt Valley, MD) and stained with 1:2000 LIVE/DEAD Far Red Dead Cell Stain (LDR) (Thermo Fisher Scientific, Waltham, MA). Cells were then fixed with 3.7% formaldehyde (Sigma Aldrich, St. Louis, MO) for 30 minutes and permeabilized with 0.5% Triton X-100 in PBS. The EdU was labeled with cy3-azide (Lumiprobe, Hunt Valley, MD) for 30 min. The cells were then blocked for one hour with Odyssey blocking buffer (LI-COR, Lincoln, NE), and stained overnight at 4°C with 2 µg/ml Hoechst 33342 (Sigma Aldrich, St. Louis, MO) and a 1:1000 dilution of anti-phospho-histone H3 (pHH3) Alexa 488 (Ser10, clone D2C8) conjugated antibody (Cell Signaling Technologies, Danvers, MA). Fixed cells were imaged with a 10x objective using an IXM-C microscope and analyzed using MetaXpress software (Molecular Devices, San Jose, CA). Nuclei were segmented based on their Hoechst signal. DNA content, defined by the total Hoechst intensity within the nuclear mask, was used to identify cells in the G1 and G2 phases of the cell cycle. The average LDR, EdU and phospho-histone H3 intensities within the nuclear masks were determined and used to classify cells as dead, in S phase or in M phase respectively. Cells with intermediate DNA content and no EdU signal were classified as S phase dropout cells. Live cell counts were normalized to DMSO-treated controls on the same plates to yield normalized growth rate inhibition (GR) values as described previously [30].

### Autophagy

RPE1 cells (1500 cells/well in 50 µl DMEM/F12, 10 % FBS, and 1 % P/S) stably expressing the GFP-LC3-RFP-LC3ΔC autophagy flux reporter [35] were seeded in 384-well plates and grown for approximately 18 h. 50 µl media with 2 x compound concentration was added and plates were subsequently placed and monitored in an IncuCyte® (Sartorius) instrument. Cells were scanned at indicated times for phase contrast and fluorescence intensity (GFP and RFP) to obtain information about confluence and autophagic flux, respectively. Hits are defined as compounds showing > 20% aberration of the GFP/RFP ratio compared to the control compounds DMSO (0.1 %) and Torin1 (250 nM) in at least 5 or more consecutive time points (equivalent to 10h screening time). Compounds were tested in triplicate and the complete screen was performed twice.

## Supporting information

S1 File. KCGS Material Trust Agreement

S2 File. Protocol for dilution and use of KCGS

S5 Table. Plating densities and media

S7 Table. Autophagy raw data

S3 Table. SMARTS analysis

S4 Table. Cell toxicity screen

S2 Table. KCGS kinase coverage

S1 Table. KCGS kinase profiles

S6 Table. GR values

## Supporting information

**S1 File. KCGS Material Trust Agreement**

(PDF)

**S2 File. Protocol for dilution and use of KCGS**

(PDF)

**S1 Table. KCGS kinase profiles**

(XLSX)

**S2 Table. KCGS kinase coverage**

(XLSX)

**S3 Table. SMARTS analysis**

(XLSX)

**S4 Table. Cell toxicity screen**

(XLSX)

**S5 Table. Plating densities and media**

(XLSX)

**S6 Table. GR values**

(XLSX)

**S7 Table. Autophagy raw data**

(XLSX)

