## Supplementary material for "The Kinase Chemogenomic Set (KCGS): An open science resource for kinase vulnerability identification": S1 File. KCGS Material Trust Agreement

By accepting the Material, you agree to hold the Material in trust for the benefit of the Beneficiaries, and, in your capacity as a trustee, you agree to the following obligations and responsibilities with respect to the Material:

1. You will not file an application for patent, nor actively seek any other form of intellectual property protection, claiming or covering the Material or its physical forms, methods of synthesis, uses, formulations, or dosages, nor will you permit others under your direct supervision to seek, nor will you assist others in seeking, any such patents or other intellectual property protections. If any invention or discovery pertaining to the Material, whether patentable or not, is first conceived or reduced to practice by you, your Establishment, or those under your direct supervision, during research using the Material, you agree not to enforce any rights that you may have in the invention or discovery against the SGC or its members, and you hereby grant to the SGC and its members a non-exclusive, irrevocable, perpetual, world-wide, sub-licensable, and royalty-free research use license under any such rights.

2. You agree to place the results of the research performed with the Material by you, your Establishment, and those under your direct supervision, in the public domain, by publishing results in one or more open access journals and by depositing the underlying data in a public access repository or, if not feasible, in an institutional repository accessible to academic researchers, within one year of publication.

3. You agree that you, your Establishment, and those under your direct supervision will acknowledge the providers listed in Appendix A as the source of the Material in all publications resulting from, and in all presentations in scientific fora regarding, use of the Materials.

4. You agree to provide the SGC with a brief update on your progress with the Materials and a report briefly describing the results of the research undertaken with the Material, upon conclusion of such research. You agree to provide a copy of all proposed publications to the SGC prior to submission.

5. You agree to have your name and institution listed as a recipient of KCGS on the SGC website and SGC publications.

6. You agree that the Material is for in vitro laboratory use only and will not be used in humans or animals, without first obtaining written consent from SGC.

7. You agree to use the Material in compliance with all applicable Federal, Provincial, State, Country, and local laws, regulations, and ordinances.

8. You agree that the Material will be used for teaching or not-for-profit research purposes only.

9. You agree not to distribute the Material to, or allow access to the Material by, any third parties without the written consent of SGC.

10. In situations where the EU GDPR applies, I understand and agree that by checking the box and pressing the Submit button that I am authorizing this site and UNC Chapel Hill to collect and process personal information about me for the purpose of interacting with this site and for the

applicable purposes listed below: Displaying a list of the recipients of KCGS on the SGC website; Receiving periodic updates from the SGC on the expansion or characterization of KCGS; Receiving requests from the SGC for input on how to improve KCGS. In the case where the EU GDPR applies and I wish to withdraw my consent for my personal information to be retained by this site or UNC-Chapel Hill, I understand that I will need to contact the site administrator and UNC-Chapel Hill. My consent to the above processing is informed, freely given, for the specific purposes listed herein, and represents an unambiguous indication of my wishes to agree to the processing of personal data related to me.
