## Supplementary material for "The Kinase Chemogenomic Set (KCGS): An open science resource for kinase vulnerability identification": S2 File. Protocol for dilution and use of KCGS

96-well plate assay single concentration screen at 1 μM

Step 1: Add to each well of the KCGS plate 9 μL of DMSO (initial concentration 10 mM; final concentration 1 mM)

Step 2: Take 1 μL from each well and transfer to a daughter plate in the desired screening format

Step 3: Add 9 μL of media to each well of the daughter plate (initial concentration 1 mM; final concentration 100 μM)

Step 4: Add 1 μL of the liquid from Step 3 to the cell-based assay (100 μL total assay volume) to generate a final concentration of 1 μM of the inhibitor with a DMSO concentration of 0.1%.

These steps provide enough volume to perform an assay allowing for both technical and biological replicates as well as a counter screen in a control cell line.
